# ALTRE: workflow for defining ALTered Regulatory Elements using chromatin accessibility data

**DOI:** 10.1101/080564

**Authors:** Elizabeth Baskin, Rick Farouni, Ewy A. Mathe

**Affiliations:** Department of Biomedical Informatics, College of Medicine, The Ohio State University, Columbus, OH 43210, USA

## Abstract

**Summary:** Regulatory elements regulate gene transcription, and their location and accessibility is cell-type specific, particularly for enhancers. Mapping and comparing chromatin accessibility between different cell types may identify mechanisms involved in cellular development and disease progression. To streamline and simplify differential analysis of regulatory elements genome-wide using chromatin accessibility data, such as DNase-seq, ATAC-seq, we developed ALTRE (ALTered Regulatory Elements), an R package and associated R Shiny web app. ALTRE makes such analysis accessible to a wide range of users – from novice to practiced computational biologists.

**Availability:** https://github.com/Mathelab/ALTRE

## 1 Introduction

Assays that measure chromatin accessibility genome-wide, such as FAIRE-seq (Giresi *et al.*, 2007), DNase-seq (Crawford *et al.*, 2006; John *et al.*, 2013; Thurman *et al.*, 2012), and ATAC-seq (Buenrostro *et al.*, 2013), enable global mapping of regulatory elements (REs), including promoters and enhancers. Organization of these REs, particularly enhancers, is cell-type specific (Kieffer-Kwon *et al.*, 2013; Rendeiro *et al.*, 2016; Stergachis *et al.*, 2013) and is a strong determinant of disease mutational landscapes, including those of cancer (Polak *et al.*, 2015). Thus, identifying REs that differ in accessibility between cell types, such as cancerous and noncancerous cell lines and tissues, holds promise for pinpointing mechanisms involved in disease progression. Furthermore, REs that control disease-related genes and pathways can be investigated as putative therapeutic targets, or may even be such targets themselves (Heinz *et al.*, 2015; Lam *et al.*, 2013).

To the best of our knowledge, no comprehensive and user-friendly workflow for downstream analysis of chromatin accessibility data is available. Downstream analysis includes guiding chromatin accessibility alignment and peak data to interpretable results of REs and pathways of interest. However, there are no standardized approaches or guidelines. Typically, individual data analyses pipelines must be created from scratch in-house, thereby making reproducible, shareable data-analysis difficult. ALTRE provides a workflow so users can identify altered REs between two different cell types or conditions, and includes a Shiny (RStudio shiny: Easy web applications in R. 2014) web interface for those not as fluent in the R statistical language.

## 2 Implementation

### 2.1 Data preparation and set-up

Typical of high-throughput sequencing data, chromatin accessibility data are delivered in FASTQ files. Quality control, alignment, and peak calling of the FASTQ file reads, described in detail elsewhere (Baek *et al.*, 2012; Boyle *et al.*, 2008; Jalili *et al.*, 2016; Rashid *et al.*, 2011; Zhang *et al.*, 2008), must be performed before using ALTRE. To start the ALTRE workflow, users need to generate a comma-separated-values CSV file with 4 columns for each sample to be analyzed: 1) name of alignment (BAM) files; 2) name of peak (BED) files; 3) sample name; 4) replicate number. All files should be placed in the same folder and the software will detect the location of the files when reading in the CSV. A minimum of 2 replicates per sample is required to run the workflow. To get started with ALTRE, users need to have R (≥ 3.2.0) installed.

### 2.2 General aspects and design

ALTRE was designed to be user-friendly and to streamline differential analysis of REs genome-wide. The steps of the workflow analysis are delineated in Figure 1 and include loading data, defining consensus peaks (found in multiple replicates), annotating (e.g. Transcription Start Site (TSS)-distal and TSS-proximal) and optionally merging peaks, identifying significantly altered REs based on quantitative data using DESeq2 (Love, et al., 2014), creating tracks for visualizing categorized REs in a genome browser, comparing altered REs with those defined based on binary (peak present/absent) data only, and finally, defining pathways that are enriched in cell- or condition-type specific or shared REs using GREAT (McLean *et al.*, 2010).

**Fig. 1.**
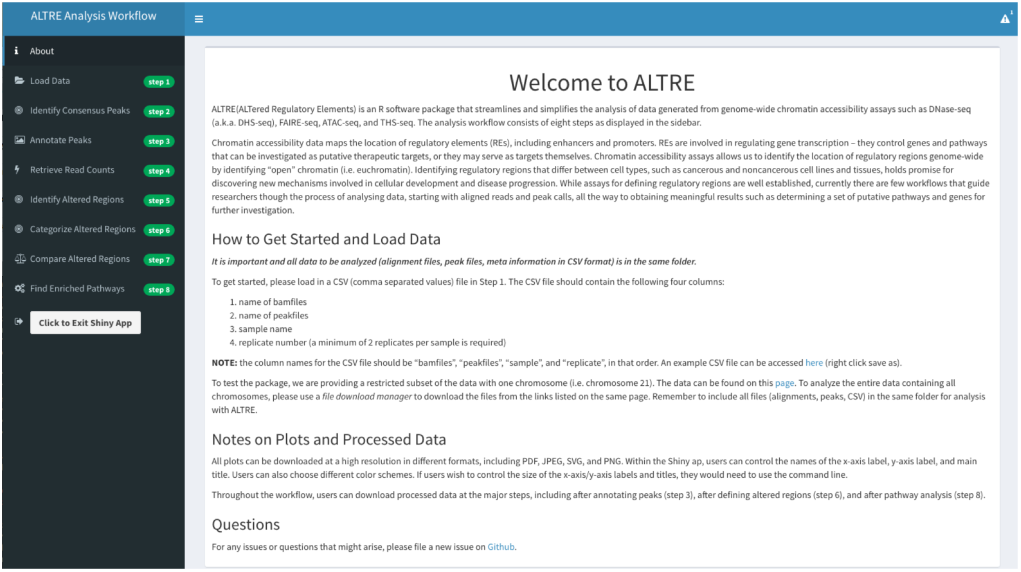
Front page of ALTRE Shiny web application showing workflow steps.

ALTRE’s embedded Shiny app takes alignment files (BAM format) and hotspot/peak files (BED format) as input. The workflow guides users through the steps described above and delineated in Figure 1. At each step, users can define thresholds, such as number of replicate samples required to define a peak as consensus, and fold changes and p-value cutoffs for definition of cell type specific or shared REs. Users can then quickly retrieve summary statistics and visualization plots (heatmaps, barplots) to ensure the appropriateness of their parameters. For ease of use, default options are provided at each step for guidance. Of note, while tools for differential binding and annotation of sequencing data exist (Bailey *et al.*, 2013; Chabbert *et al.*, 2016; Ross-innes *et al.*, 2012; Yu *et al.*, 2015; Zhu, 2013; Zhu *et al.*, 2010)(Stark and Brown, ‘DiffBind: differential binding analysis of ChIP-Seq peak data’ 2011), ALTRE supports peak merging and annotation, differential analysis, and pathway enrichment analysis in one streamlined tool.

## 3 Results and Discussion

Users can install ALTRE with the function install_github() from the devtools R package (Wickham H and Chang, W. 2016. devtools: Tools to Make Developing R Packages Easier). Full installation instructions are found at https://github.com/Mathelab/ALTRE. Users can then run the workflow either in the R console or by launching the embedded web application by typing “runShinyApp()” in the R console. A detailed vignette (https://mathelab.github.io/ALTRE/vignette.html) walks users through an example workflow analysis step-by-step.

A sample dataset is provided on GitHub and can be accessed at https://mathelab.github.io/ALTREsampledata/. This sample dataset includes ENCODE data for cancerous and associated non-cancer lung cell lines, A549 and SAEC, respectively. On a machine with 16 GB memory and a 2.5 GHz Intel Core i7 processor, the workflow takes ~ 334 seconds to complete for the example dataset using all chromosomes.

For real-time analysis of results, the ALTRE Shiny app enables users to change their parameters and directly visualize the effect of those changes through summary statistics tables and plots. For example, users can readily visualize the number of REs that are sample-type specific or shared based on their input fold change and adjusted p-value thresholds through a volcano plot and an associated statistics table. In addition, processed data can be saved after key steps in the analysis and all plots can be modified (e.g. colors) and saved as high resolution images.

With the increasing interest in researching REs to better understand transcriptional regulation and diseases, and improvements in techniques to assess these regions (Buenrostro *et al.*, 2013), chromatin accessibility assays are being increasingly generated. With this in mind, ALTRE provides a user-friendly workflow that guides the analysis and interpretation of these data.

## Acknowledgements

This work was supported by the Ohio State University Translational Data Analytics. Conflict of Interest: none declared.

## Supplementary Material

### Genome-wide (re)programming in cell differentiation and disease

Genome-wide programming of the chromatin landscape is a mechanism by which cells are differentiated into a variety of cell types that make up the human body. Changes in the chromatin structure enable accessibility and activation of regulatory regions, upon which transcription factors bind, leading to the activation of genes and associated molecular functions. Many diseases of cellular dysfunction, such as cancer, are accompanied by a similar transcriptional reprogramming, but with much less favorable consequences. Useful functions of the cell are lost and damaging functions, such as increased cellular proliferation, are gained. Recently, strong ties between the distribution of cancer-causing mutations and the regulatory landscape of the cancer’s tissue of origin have been uncovered (Polak *et al.*, 2015), a finding which positions the regulatory landscape as a prime suspect in cancer progression. Assays such as FAIRE-seq (Giresi *et al.*, 2007), DNase-seq (Crawford *et al.*, 2006; John *et al.*, 2013; Thurman *et al.*, 2012), and ATAC-seq (Buenrostro *et al.*, 2013) can investigate these changes in chromatin structure genome-wide by capturing the location of open, or transcriptionally active, chromatin (Kieffer-Kwon *et al.*, 2013; Stergachis *et al.*, 2013). With these assays, peaks signal the presence of open chromatin regions, which mark the location of regulatory elements (REs), such as promoters and enhancers, that control gene expression. Identifying REs that differ in accessibility between cell types, such as cancerous and noncancerous cell lines and tissues, holds promise for identifying new mechanisms involved in cancer progression. These regions control the genes and pathways that can be investigated as putative therapeutic targets, or may even be such targets themselves.

### The problems in data analysis ALTRE solves

Currently, no comprehensive and user-friendly workflow for downstream analysis of chromatin accessibility data, from aligned reads and peaks to meaningful results of genes and pathways of interest, exists. Instead, individual data analyses pipelines must be created from scratch in-house, making reproducible, shareable data-analysis difficult, as there are no standardized approaches. Additionally, the burden of both time investment and programming skill is a deterrent to individuals new to the field, especially those with little computer programming background. Thus, the scientific community will benefit from a user-friendly software tool that 1) guides newcomers to an understanding of the analysis process, 2) offers a streamlined, tested tool for the analysis of chromatin accessibility data, and 3) delivers meaningful results, including pathway enrichment analysis, that can be leveraged for further studies.

### General aspects and design

ALTRE is a workflow package that takes alignment (BAM format) and peak (BED format) files as input. The workflow guides users through visualizing areas of open chromatin as peaks in a genome browser, defining consensus peaks (found in multiple replicates), annotating peaks into candidate TSS-proximal and TSS-distal regions (based on their proximity to known Transcription Start Sites), identifying significantly altered TSS-proximal and TSS-distal regions based on accessibility, and identifying enriched pathways in cell-type specific or shared REs using GREAT (McLean *et al.*, 2010). The dysfunction of these pathways may play a role in cell type specificity or disease progression, if diseased cells are compared to normal cells. Of note, the location of REs relies on peak caller algorithms, which differ in their sensitivity of peak calls (e.g. due to different peak shapes or signal to noise ratios), even between replicate samples (Koohy *et al.*, 2014). While one can use biological replicates to call consensus (e.g. reproducible) peaks (Jalili *et al.*, 2016), ALTRE will identify consensus peaks after peak calling that are defined among multiple replicate samples. In addition, ALTRE defines altered peaks using both binary data (e.g. presence/absence of peaks) and quantitative data (e.g. peak intensity) and enables the comparison of results from both methods. For quantitative analysis, counts in REs and the DESeq2 algorithm (Love *et al.*, 2014) are leveraged. While tools for differential binding and annotation of sequencing data exist (Bailey *et al.*, 2013; Chabbert *et al.*, 2016; Ross-innes *et al.*, 2012; Yu *et al.*, 2015; Zhu, 2013; Zhu *et al.*, 2010), ALTRE supports peak merging and annotation, differential analysis, and pathway enrichment analysis in one streamlined analysis tool.

### Software access and installation instructions

ALTRE is hosted on GitHub (http://mathelab.github.io/ALTRE), allowing developers and users to readily see, use, and modify the code. The vignette for the software is available at https://mathelab.github.io/ALTRE/vignette.html, and the code that generates the vignette is accessible at https://github.com/Mathelab/ALTRE/tree/gh-pages. Users can install the package with the function install_github() in the devtools package (Wickham H.; Chang, W., devtools: Tools to Make Developing R Packages Easier. 2016), thereby greatly simplifying the R package set-up. The following lines of code will install the package and all its dependencies:

~~~
source(“http://bioconductor.org/biocLite.R”)
BiocInstaller::biocLite(c(‘org.Hs.eg.db’,
                          ‘EnsDb.Hsapiens.v75’,
                          ‘GO.db’))
install.packages(“devtools”)
devtools::install_github(“mathelab/ALTRE”)
~~~

The project’s GitHub page (https://github.com/Mathelab/ALTRE) contains detailed information on installation and on troubleshooting installation errors (e.g. XML package installation error on Linux systems). If errors occur (as could happen with older systems), users can submit a bug report on the issues page of the project’s GitHub site and we will promptly respond to the issues. Finally, we incorporated Travis-CI, a continuous integration service used to build and test software projects so that we can automatically test if ALTRE successfully builds and installs after any modification to the code.

### Example data

Our example data includes processed sequencing reads from ENCODE DNase-seq experiments. This data was measured in lung cancer cell line, A549, as one group, and the corresponding normal cell line, SAEC, as “reference”. A restricted subset of this data, comprising only chromosome 21, can be found at http://mathelab.github.io/ALTREsampledata/. The CSV file that is input into the first step of ALTRE, function loadCSVFile(), can be downloaded separately at https://raw.githubusercontent.com/mathelab/ALTREsampledata/master/DNaseEncodeExample.csv. All downloaded files (alignment files, peak files, and CSV) should be placed in the same folder before using ALTRE. The corresponding entire datasets can be downloaded directly from the ENCODE project. The links for the sample datasets are are provided on the website as well.

### Workflow

The workflow comprises 8 steps, where the first step processes the CSV sample information file. The folder with the CSV file must also contain all analysis files, including alignment (BAM format) and peak files (BED format). The outputs of the major functions (Steps 2-7) applied to our sample data are shown and discussed below. Of note, if the Shiny web application is used by running runShinyApp() in the R console), then users can save data at the major steps (Steps 3, 6, 7, and 8 in the web interface). In the R console and web application, all plots can be modified (e.g. colors, main title and axis text) and saved as high resolution images. In the R console, the size of the main title and axis text can also be modified.

#### Defining consensus peaks (Step 2)

ALTRE analysis requires two different types of samples and at least two biological replicates of each cell type. Consensus peaks are those present in at least N replicates, where N is input by the user. The first step in the worflow takes an input of two or more peak files per sample and returns a genomic ranges object representing consensus peaks. Additionally, the function will output metrics summarizing how many peaks were present in each replicate for each sample type, and how many peaks are “consensus”, or present in at least N replicates (Figure 1).

**Fig. 1.**
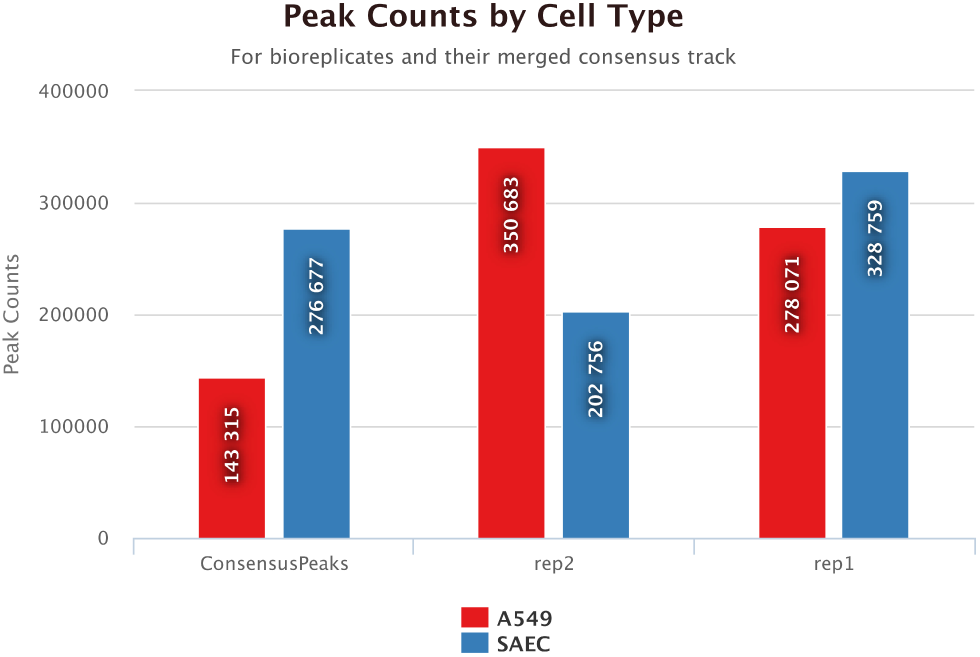
Consensus peaks boxplot (Step 2)

The function plotConsensusPeaks() produces the plot in Figure 1.

#### Annotation of Regulatory Elements (Step 3)

Consensus REs are categorized as either candidate TSS-proximal or TSS-distal. By default, REs within 1,500 bp of a Transcription Start Site (TSS) are considered TSS-proximal, while those beyond 1,500 bp of a TSS are considered TSS-distal. This distance can be modified by user input. Also by default, the location of the TSS are derived from Ensembl, through the R package org.Hs.eg.db, Carlson M. org.Hs.eg.db: Genome wide annotation for Human. R package version 3.2.3.

It is reasonable to think that REs within a small arbitrary distance (e.g. 1000-1500 bp) belong to the same region and should be merged (the accuracy of peak boundaries is questionable). This merging option can be implemented by setting the “merge” argument to TRUE. Furthermore, the merging can be implemented within RE type (i.e. only TSS-distal regions will be merged with other TSS-distal regions, and likewise, TSS-proximal only with TSS-proximal), or without regard to RE type by setting the arguments “regionspecific”, “distancefromTSSdist”, and “distancefromTSSprox” accordingly.

The function plotCombineAnnotatePeaks() produces the plot in Figure 2. Users can also download the annotated regions by calling the function writeConsensusRE().

**Fig. 2.**
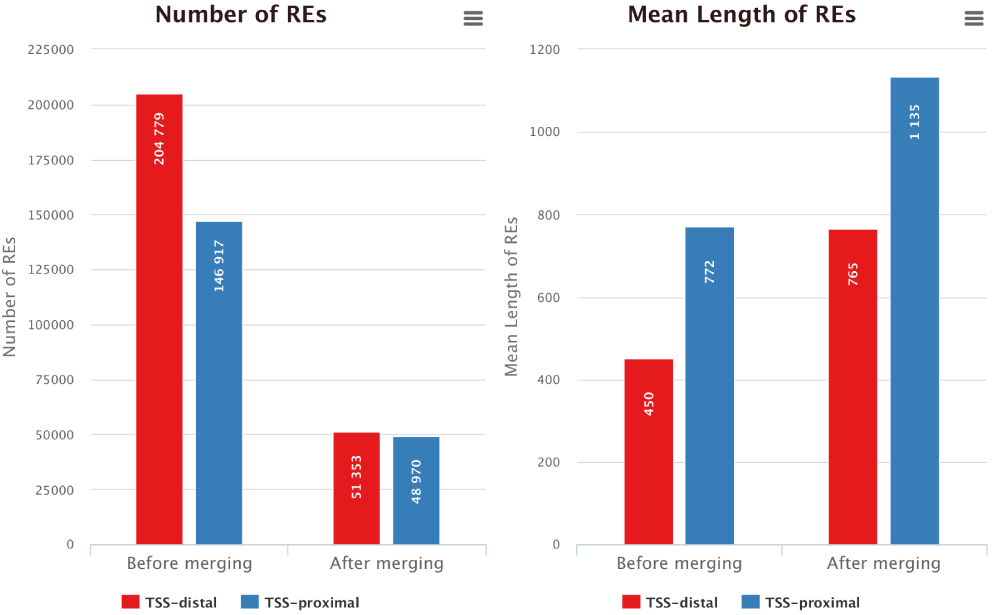
Number of candidate TSS-proximal or TSS-distal REs after annotation (Step 3)

#### Identification of altered REs (Steps 4-6)

With consensus REs identified and categorized for each cell type, the most integral aspect of the pipeline can be started: identification of cell-type specific and shared REs based on peak intensity, which measures the extent of chromatin accessibility. First, the number of reads, which approximates the accessibility, in each RE must be retrieved with the function getCounts(). Second, the DESeq2 algorithm for count-based differential analysis is implemented by calling the function countanalysis(), which takes the log2 fold change and p-value cut-off parameters used by the DESeq2 results() function to optimize hypothesis testing and calculation of adjusted p-values, respectively. See https://www.bioconductor.org/packages/devel/bioc/vignettes/DESeq2/inst/doc/DESeq2.pdf for more details.

The third function, categAltrePeaks() applies user input criteria to categorize REs as experiment-specific (higher accessibility in experiment sample), reference-specific (higher accessibility in reference sample), or shared. Input criteria include: 1) lfctypespecific, pvaltypespecific: log2 fold change and p-value cutoff for categorizing experiment- or reference-specific REs; and 2) lfcshared, pvalshared: log2 fold change and p-value cutoff for categorizing shared REs (those with log2 fold changes within lfcshared and with adjusted p-values > pvalshared).

The functions plotCountAnalysis() and plotDistCountAnalysis() produce the plots in Figure 3.

**Fig. 3.**
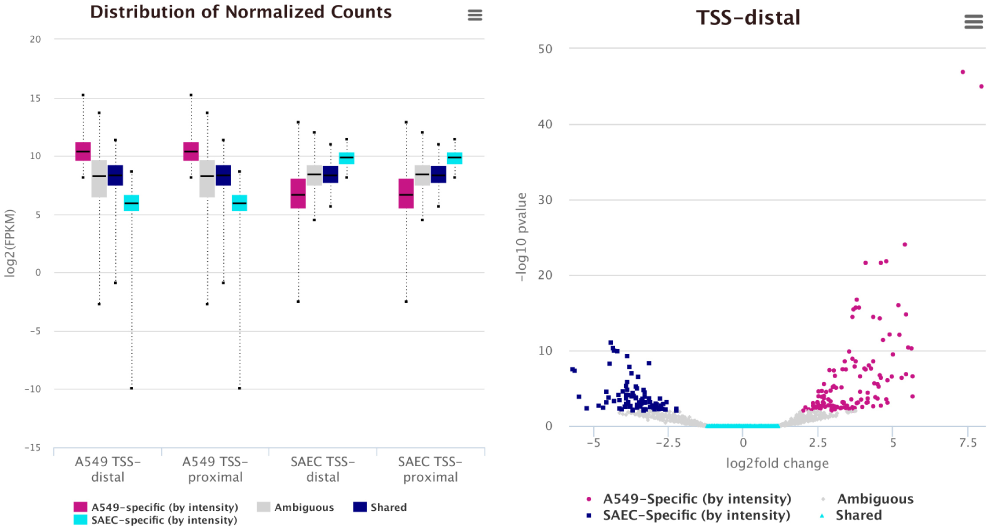
Distirubtion of intensities for altered REs (left) and volcano plot (right) (Steps 4-6)

Additionally, the function writeBedFile() generates BED files to display peaks that are color-coded by cell type specificity according to the categAltrePeaks() output. Viewing these track files along with the raw data allows visual inspection of regions for consistency. For example, a red, type-specific region should have a tall peak in one cell type and little to no peak in the other, while a purple, shared region should have comparable peaks in both cell types.

#### Comparing REs defined from binary vs quantitative data (Step 7)

ALTRE uses peak intensity as a proxy for the amount of chromatin accessibility, to categorize REs as cell type specific or shared. Another method for defining cell type specificity of REs is to use peak/binary information (whether or not the RE location has a peak).

The function comparePeaksAltre() compares results obtained when the quantitative/peak intensity data is used, and when the peak/binary information is used. The plotCompareMethodsAll() function displays the results in Figure 4. As expected, leveraging the peak intensity reduces the number of REs categorized, mainly because “ambiguous” peaks that do not meet the shared or cell-type specific criteria are no longer categorized.

**Fig. 4.**
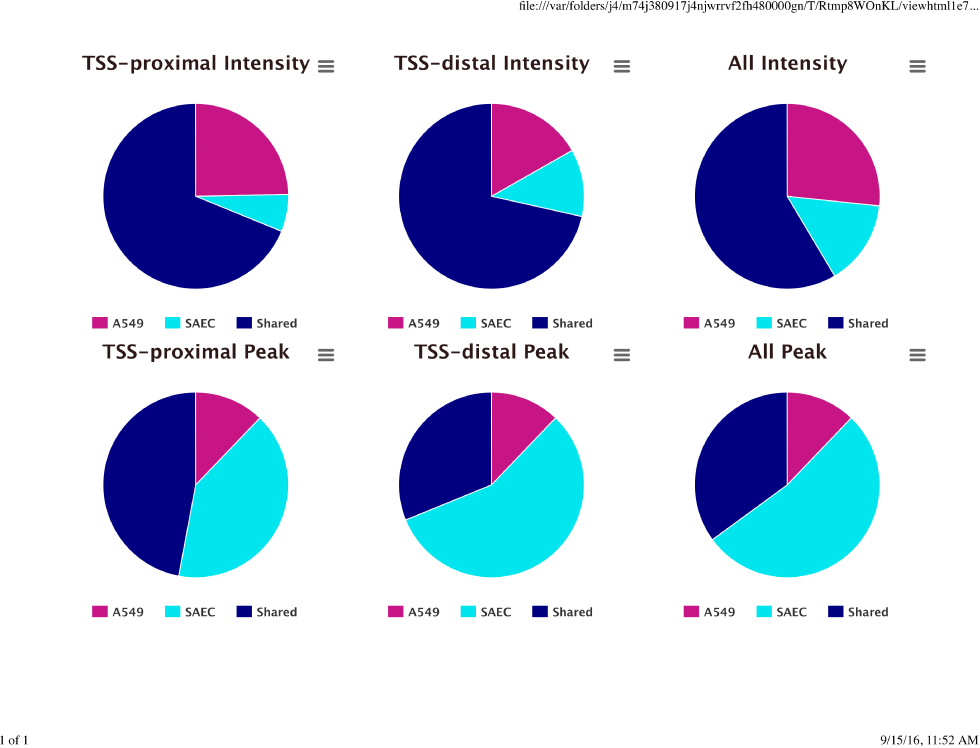
Comparing categorization based on binary or quantitative data (Step 7)

Users can write the results of CompareMethodsAll() with the function writeCompareRE(). The function will create a CSV file that denotes the number of peaks called as experiment - or reference-specific and shared using the quantitative/peak intensity or peak/binary approach.

#### Pathway enrichment of genes closest to altered REs (Step 8)

The runGREAT() and processPathways() functions perform pathway enrichment analysis on REs categorized as experiment-specific, reference-specific, and shared. These functions rely on the GREAT algorithm (McLean, et al., 2010), which incorporates distal sites when mapping REs to genes, and calculates pathway enrichment using binomial testing, which controls false positives based on the annotation of genes that are nearby REs of interest. An R interface to the GREAT algorithm (Gu, Z, 2016. rGREAT: Client for GREAT Analysis) is embedded into ALTRE. By default, the Gene Ontology (Gene Ontology, 2015) pathway (annotations include molecular functions, biological processes, and cell components) annotation is used. The function returns a list of enriched pathways for each type of RE (e.g. experiment- and reference-specific, shared), according to user-input p-value and fold change cutoff.

The function plotGREATenrich() produces the plot in (Figure 5). Note that this plotting function will only plot the top 10 pathways for each category by default. To view all the results, change the default (numshow category) or write all the results by calling the function writeGREATpath().

**Fig. 5.**
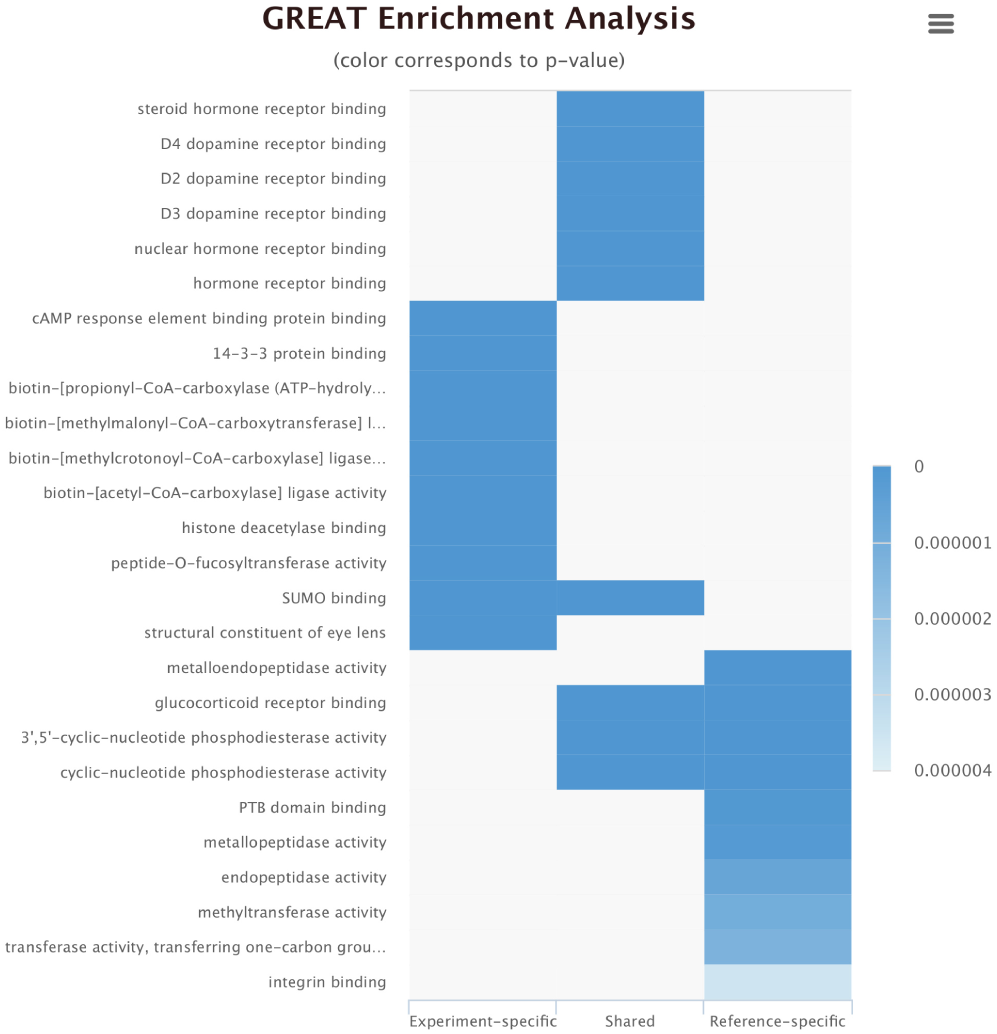
Pathway enrichment (Step 8)

## Acknowledgements

This work was supported by the Ohio State University Translational Data Analytics.

